# Principles of resilient coding for plant ecophysiologists

**DOI:** 10.1101/2020.09.11.293530

**Authors:** Joseph R Stinziano, Cassaundra Roback, Demi Gamble, Bridget K Murphy, Patrick J Hudson, Christopher D Muir

## Abstract

Plant ecophysiology is founded on a rich body of physical and chemical theory, but it is challenging to connect theory with data in unambiguous, analytically rigorous, and reproducible ways. Custom scripts written in computer programming languages (coding) enable plant ecophysiologists to model plant processes and fit models to data reproducibly using advanced statistical techniques. Since many ecophysiologists lack formal programming education, we have yet to adopt a unified set of coding principles and standards that could make coding easier to learn, use, and modify. We identify eight principles to help in plant ecophysiologists without much programming experience to write resilient code: 1) standardized nomenclature, 2) consistency in style, 3) increased modularity/extensibility for easier editing and understanding, 4) code scalability for application to large datasets, 5) documented contingencies for code maintenance, 6) documentation to facilitate user understanding; 7) extensive tutorials, and 8) unit testing. We illustrate these principles using a new R package, {photosynthesis}, which provides a set of analytical and simulation tools for plant ecophysiology. Our goal with these principles is to advance scientific discovery in plant ecophysiology by making it easier to use code for simulation and data analysis, reproduce results, and rapidly incorporate new biological understanding and analytical tools.

## Background

Computer coding is becoming an increasingly important skill in biological research (Sayres *et al.*, 2018), especially within plant ecophysiology. A disconnect in coding skill and a lack of formal computer science training can make it difficult for biologists to create or modify programs to incorporate new understanding of biological processes. In other words, sophisticated code (by trained programmers) is efficient, but difficult to modify by biologists for new uses. So why code at all? Coding allows for consistent, reproducible, transparent, and scalable analyses of scientific data, while at the same time minimizing human workhours compared to using pre-packaged software. However, most published ecophysiological analyses use spreadsheet-based methods rather than computer code, which comes with some limitations. For example, Sharkey *et al.* (2007) have an Excel spreadsheet-based method for fitting photosynthetic CO_2_ response (*A-C_i_*) curves (also see Bellasio *et al.*, 2016). A spreadsheet-based method can take several minutes per curve and involves a substantial amount of subjective decision-making (e.g. ‘eye-balling’ where transitions between CO_2_- and RuBP-limited photosynthesis occur). Likewise, analysis of pressure-volume curves for hydraulic parameters is usually done via an Excel spreadsheet-based method (Sack *et al.*, 2003), which can be time-consuming, requires subjective decisions, and spreadsheets are usually not published with manuscripts, obscuring methodology. The total workload is time per spreadsheet multiplied by the number of curves, which can be inefficient in large studies. Cryptic changes in the spreadsheets can occur without a record of the change, potentially leading to compounding errors. Furthermore, spreadsheet tools often break, requiring a fresh, unaltered spreadsheet to be used for each CO_2_ response curve. Another option, provided by Gu *et al.* (2010) (leafweb.org) provides an online service that analyses *A-C_i_* curves, however in this case, the analysis is a black-box and could be misused by users lacking an understanding of the fitting process, and the data are stored on a government server which may cause some users discomfort.

Meanwhile, Duursma (2015) developed an R package, {plantecophys}, that can obtain similar outputs to the Sharkey et al. (2007) fitting tools in seconds, with far fewer subjective decisions that can easily be outlined in the code used in the fitting process, while providing a similar, but transparent approach as in Gu et al. (2010). Like the {plantecophys} package, analytical methods should be fully transparent and reproducible. As such, authors should publish their code, which is still not the norm in plant ecophysiology (but see Kumarathunge et al., 2019 for an example of published code). As a community, increased adoption and dissemination of code will help the field perform more sophisticated analyses and model comparison (e.g. Walker *et al.* 2021). Coding may also streamline integration between theory and data analysis, especially for complex mathematical formulations that require computationally intensive numerical methods, a common situation in plant ecophysiology. Ideally, we would like a workflow in which we state our assumptions mathematically, derive empirical predictions, and test those predictions or estimate parameters with data. The process of translating a mathematical model of biology into code can also help novice and advanced coders deepen their understanding of models and their assumptions before confronting them with data. Open-source, research-grade computer algebra systems like SymPy (Meuer *et al.*, 2017) and numerical solvers aid mathematical derivation and are part of or can be readily integrated with programming languages that are widely used for data manipulation and analysis, such as R (R Core Team, 2021), Python (Python Software Foundation), or Julia (Bezanson *et al.* 2017).

Although coding can speed up large analyses, reduce errors, make analyses reproducible, and integrate theory with data, writing robust code that can be understood and reused by other scientists is not easy. First, one must learn one or more programming languages (e.g. R, Python, Matlab, Julia), which can involve steep learning curves. Second, even though coding one’s own analysis can make it easier to catch errors associated with inappropriate use of black-box proprietary software, one must still understand the assumptions and limitations of statistical techniques and conceptual tools. Finally, code can be as unique as someone’s handwriting, which can make it difficult even for an experienced programmer to make sense of a “transparent” analysis unless there is sufficient annotation within the code.

In this perspective, we propose seven principles of coding tailored to the specific needs of the plant ecophysiology research community. For example, guidance in other scientific fields often emphasize computational speed. However, given the typical scale of ecophysiological datasets (~MB, i.e. small-batch, artisanal datasets) and the computer power of personal computers (~GB of RAM, ~GHz of processing power), computational speed is usually not a major limitation. Instead, ecophysiologists often need to estimate parameters derived from complex biophysical/chemical models. Coding is important as the complex models required to fit different response curves involve many interacting equations, numerical solvers, and parameters that either need to be set or estimated. For example, there are seven different models that can be used to fit temperature responses which ultimately require different equations and fixed parameters (Arrhenius, 1915; Johnson *et al.* 1942; Medlyn *et al.*, 2002; Kruse *et al.*, 2006; Heskel *et al.*, 2016; Liang *et al.*, 2018). In this domain, code flexibility and modularity are usually more important than computational speed. Furthermore, flexibility and modularity in code would enhance the sustainability of software after publication, which can be an issue (Prlić & Procter, 2012). Here we demonstrate coding principles designed for plant ecophysiology using a new R package called {photosynthesis}. We caution that this software is a work-in-progress that does not yet completely adhere to all of the coding principles to which we aspire, but will be refined in future releases. This perspective, written by trained biologists not programmers, is intended to convey some of the lessons we have learned so far to provide guidance for plant ecophysiologists who are thinking about or starting to code their workflows, especially using R. We recognize that many other scientists in this field are adept coders who have already honed their practices through experience. Hence, this perspective is intended to guide for less experienced coders rather than a mandate for the entire field. We hope our perspective spurs experienced coders to share “best practices” with less-experienced peers and expand the principles below to other languages besides R. As computational plant ecophysiology matures, we hope that this perspective will help move the field toward more standardized and sustainable software practices like those in more computationally-intensive subfields of biology like population genetics (Adrion *et al.* 2020).

## Description

### Principles of Coding

The overarching concept we propose is making code resilient by making it easier to use, reproduce, and modify. Obviously not every possible discovery and need within a scientific field can be predicted, but the code can be written to allow easy modification and accommodation of the source code as the science progresses. Functional programming in R and other languages provides a powerful tool for writing functions that take functions as arguments and easily process newly written code into a standardized output without the need for ever modifying the original function itself (Wickham, 2019). Such an approach helps to write modular code that is easy to modify and understand, while minimizing interdependencies between functions.

Freely available resources already exist for good coding practices in R packages and can be applied to R scripts as well, primarily from the efforts of Hadley Wickham (Wickham, 2014, 2015, 2016, 2017, 2019; Wickham & Grolemund, 2016). As well, guides to best practices for scientific computing exist (see Wilson et al. 2014 for a list of best practices). Here we propose principles of coding for plant ecophysiology that, if implemented, could circumvent some of the common coding issues encountered when modifying the code of others, reduce the learning difficulty for nascent coders, and make software maintenance much easier:

1. Standardized nomenclature for variables and functions
2. Consistent style
3. Modularity and extensibility
4. Scalability
5. Documented contingencies
6. Documentation
7. Extensive tutorials
8. Unit testing

We think that adopting some or all of these principles will improve code reproducibility and help advance scientific discovery, but our goal is not to rigidly prescribe how plant ecophysiologists should do their work. First, we recognize that others will have different, well-reasoned preferences and/or apply principles we have not covered here. Second, those who find these principles useful may find implementing all of them time-consuming at first. We strongly encourage incremental progress and not making perfection the enemy of the good. Indeed, the {photosynthesis} package described below only partially implements our principles, with much left to do in future development.

#### Principle 1: standardized nomenclature

Names vary wildly between functions with published code and data and even amongst instruments within the same company (e.g. for net CO_2_ assimilation, “*A*” is used in the Li-Cor 6800 and “PHOTO” is used in the Li-Cor 6400). Ideally, we need both standardized nomenclature in the field (e.g. Reid *et al.* 2005) and standardized construction of variable and function names to enhance readability and reduce the burden for learning how to use new packages and functions or testing published code. For example, g is always in reference to conductance, where a subscript term would then describe the physical pathway (e.g. s for stomata, c for cuticle, or m for mesophyll) as well as the gas (e.g. c for CO_2_, w for water vapor). For example, *g_sw_* would mean stomatal conductance to water vapor. Standardizing nomenclature across both mathematical models and data files can also streamline theory-data integration, but this also requires standard translation between mathematical and computer notation, which is beyond our scope here.

For example, in {photosynthesis}, every function is named in a descriptive manner: e.g. `fit_t_response` fits specified temperature responses model to data, while `fit_gs_model` fits specified models of stomatal conductance. Variable names are also standardized: e.g. “T_leaf” always means leaf temperature in Kelvin (K), “A_net” always means net CO_2_ assimilation in μmol m^-2^ s^-1^. In this regard, standard units should also be imposed in the analysis (e.g. in R via the {units} package (Pebesma et al., 2016)), to remove any ambiguities when interpreting the output. To allow for differences in variable names from the raw data (e.g. from using different machines), the “varnames’’ list is used to translate input names (note that this convention is adopted from {plantecophys} (Duursma, 2015)). We propose adopting Wickham’s (Wickham, 2019) style in that functions that *do* something have a verb name, e.g. `fit_aci_response`, while functions that act as objects within other functions (e.g. stomatal conductance models) should have a noun name, e.g. `gs_model`.

#### Principle 2: consistent style

Consistent coding style makes reading code easier - certain conventions, e.g. commenting what the *next* line of code does, can make it easier to understand code documentation. Our preference is for the ‘tidy style’, which applies to both data and code structure, and much else (see the *The tidyverse Style Guide:* https://style.tidyverse.org/). For data, tidy style advocates that each column is a variable, and each row is an observation, since R is particularly suited for this style of data structure. Popular R packages like {dplyr} (Wickham *et al.*, 2020) and {tidyr} (Wickham and Henry, 2020) facilitate tidy data and many other packages, like {photosynthesis}, use them for consistent style. For code, computers do not care about style, as long as it is correctly formatted, but for humans reading code, adherence to well-designed style can be helpful, especially for beginners trying to learn from others. A benefit of tidy style in particular is that R packages {styler} (Müller and Walthert, 2020), {lintr} (Hester *et al.* 2020), and {formatR} (Xie, 2019) can automate conformity to style. Ideally, a consistent style would be adopted across the field, however this may be too rigid. Style can be highly personal, and many experienced coders likely have developed their own style, formal or informal, that works for them. Our proposal is geared for beginning coders who are looking for guidance on an established and easy-to-implement style. At the very least, a consistent style *within* a project will make it easier to read, understand, and modify the code.

#### Principle 3: modularity and extensibility

Arguably, code written for plant ecophysiologists, whether formally trained in coding or not, should be written in a modular manner, much like Lego bricks, where one component (e.g. Arrhenius function) can be easily swapped with another (e.g. peaked Arrhenius function), or extended (e.g. hypothetical mechanistic temperature response model). Note that this may increase apparent complexity of software packages by creating more functions and make it more difficult to work with at first. However, it will make adding, subtracting, or modifying code modules easier for researchers who need to make on-the-fly changes to code as new biological processes are discovered or old ones re-evaluated. To achieve modularity in the structure of photosynthesis, we used principles of functional programming to develop a set of key functions for processing data and running quality control checks: `fit_many`, `analyze_sensitivity`, `compile_data`, and `print_graphs`. Both `fit_many` and `analyze_sensitivity` can be run with any function within and outside of {photosynthesis} to run multiple curve fits or sensitivity analyses on assumed input parameters. Meanwhile, `compile_data` is used for processing the list outputs from `fit_many` into a form usable for further analyses and export from R, and `print_graphs` is used to export all graphs from a list as either .jpeg or compiled as a .pdf.

For curve fitting functions with multiple models (e.g. temperature responses, *g_s_* models), we use a basic function (e.g. `fit_t_response`), which contains fitting procedures for each of the seven temperature response models in the package. Meanwhile, a `t_functions` file contains all the temperature response functions. To extend the capabilities and add in a new temperature response model, we simply need to add the new model to `t_functions`, and the fitting procedure to fit`_t_response`. Currently, adding new functions requires modifying the source code, but future versions should increase extensibility by allowing users to supply any temperature-response function. This principle of function building increases the extensibility of the code, while consistent style and standardized nomenclature provide the rules for writing the extended components.

Modularity also applies to modelling. The {photosynthesis} functions `photo` and `photosynthesis` model C_3_ photosynthesis using the Faquhar-von Caemmerer-Berry biochemical model (Farquhar *et al.* 1980). To account for temperature dependence, a user can specify leaf temperature, or they can provide additional inputs (e.g. air temperature, leaf size, wind speed, etc.) to model leaf temperature using energy balance in the R package {tealeaves} (Muir 2019). Both {photosynthesis} and {tealeaves} packages are modular in that they can work independently or be readily integrated (**Methods S1**). Ideally, future modeling packages would add modules to model environmental and plant parameters either on their own or integrated with these tools.

#### Principle 4: scalability

A major advantage in using code to analyze data is the ability to scale up an analysis to reduce time spent on repetitive tasks common in spreadsheet-based methods such as copy- and-paste, selecting data, choosing menu options, etc. Functions allow the same model to be fit across groups within a dataset using a consistent method. For this, our `fit_many` function and the principles of functional programming are how we achieve scalability within the package. Rather than writing functions for each type of model or curve, we have a single multiple fitting function, sensitivity analysis function, and printing function. R even has generic functions for scaling such as `apply` (base R language) and `map` ({purrr} package [Henry & Wickham, 2020]) which can be easily parallelized for speed (e.g. {parallel} and {furrr} [Vaughn & Dancho, 2018] packages). This makes it easy to scale a new function within the software to a large dataset.

#### Principle 5: documented contingencies

By documenting which functions are dependent on one another, it becomes easier to troubleshoot issues when modifying code and to pre-empt issues when adding or replacing a component. For example, `fit_aq_response` depends on `aq_response` - if we want to change from the non-rectangular hyperbola model to a rectangular hyperbolic model, then `fit_aq_response` needs to be modified in addition to `aq_response`. To document contingencies, we created a function, `check_dependencies`, which uses {pkgnet} (Burns et al., 2020) to generate an html report that automatically documents R package inter-dependencies and function inter-dependencies. This is particularly useful when adding, subtracting, or modifying functions in the package, as it allows planning to minimize issues that could break code.

#### Principle 6: documentation

Code annotations allow a new user to readily understand what a line of code is doing, how it is doing it, and why the code is written in a particular way. By providing exhaustive line-by-line annotation of a function, a new user can more rapidly understand the blueprint of the function. This is especially useful for R scripts and code hosted on GitHub (unfortunately, comments are erased from code upon submission to CRAN). For example, in `fit_t_response`, we outline the need for running looped iterations for the starting values of nonlinear least squares curve-fitting (Fig. 1). In the case of R packages hosted on CRAN, R documentation files provide information on how to use a function, though as a terser set of instructions as per CRAN policies (https://cran.r-project.org/doc/manuals/r-devel/R-exts.html).

**Figure 1.**
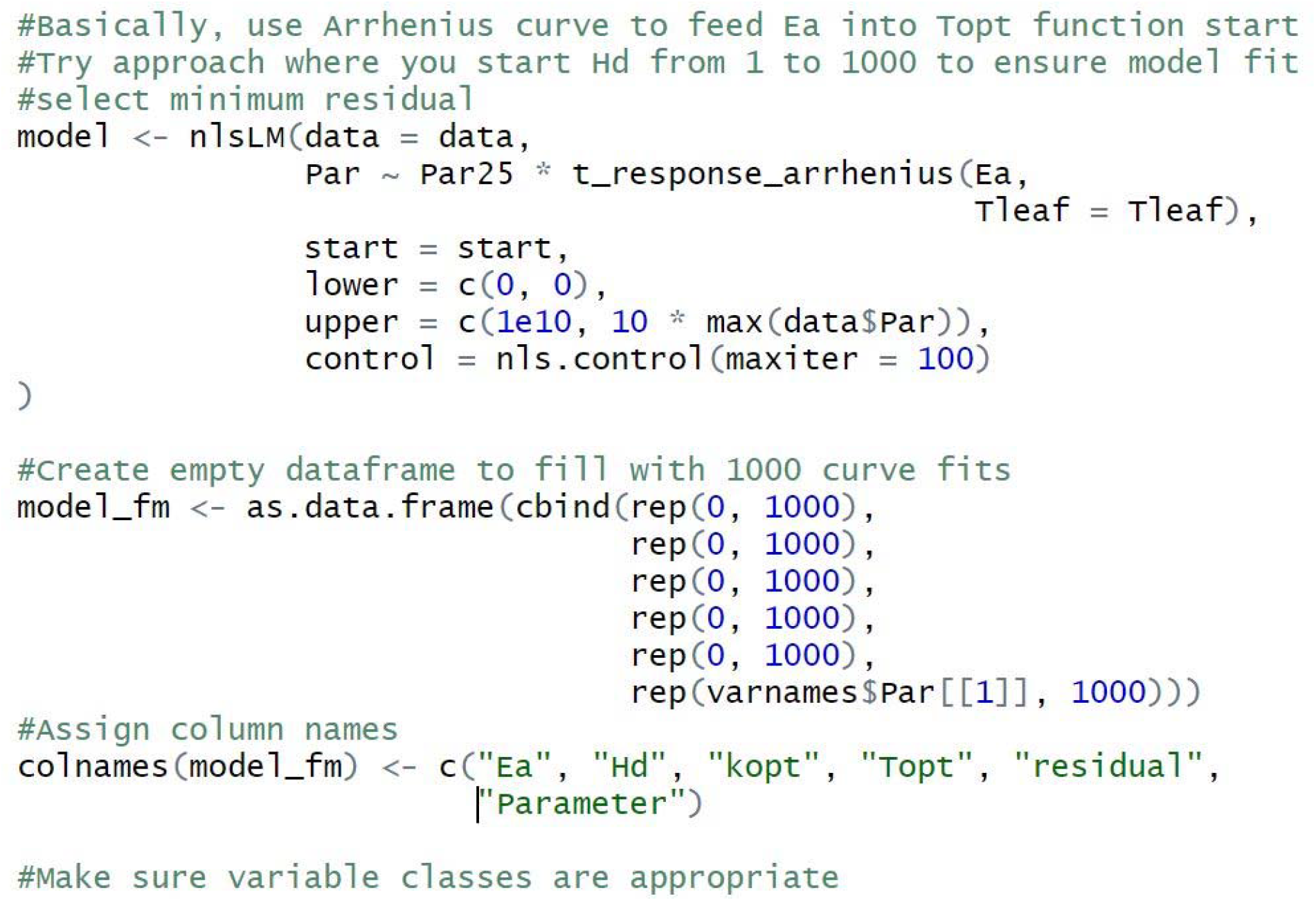
Example of coding annotations to explain the given analytical approach.

Enough metadata and commenting should be provided for a new user to understand how to use the written code (which can be an issue that affects widespread use of a program, Mangul *et al*. 2019).

#### Principle 7: extensive tutorials

As with any tool, software will only be used if potential users can understand how it works. Extensive tutorials, while providing function-by-function examples of how to use the software, should also incorporate basic data-wrangling examples and explanations of why a given approach to data analysis is used in the field. The benefits of this approach include: making the code easier to adopt into your own analysis, making it easier for new coders to learn enough of the language to use the package effectively, and help trainees learn the appropriate theory behind the measurements and analytical approach. The net effect should be to increase the inclusivity of the field by reducing barriers to success since not all individuals will have equal access to workshops or experienced colleagues.

#### Principle 8: unit testing and benchmarking

For reproducibility, code should yield the same results when it is run by other users months or years into the future. Unit testing, a common practice in software development that is still rare in scientific code, evaluates whether various components, such as custom functions, perform as expected. If all the components work as expected, it provides confidence that the whole body of code does what it is supposed to. Most scientists informally test their functions as they develop them, but formal unit testing involves writing scripts to test code and can be rerun to periodically check whether code still works as expected. More dedicated efforts automate testing and quantify code coverage, the fraction of code that is evaluated during automated tests. There are many ways to implement unit testing, but the {testthat} package is one option for R packages (Wickham, 2011) that {photosynthesis} uses for some (but not yet all) of its source code. A related concept is benchmarking, by which we mean comparing parameter estimates from the same data set using different software or later versions of the same software. Benchmarking can help determine if parameter estimation is consistent between software packages. For example, parameter estimates of photosynthetic CO_2_ response parameters (Farquhar *et al*., 1980) are very similar using comparable settings in {photosynthesis} and {plantecophys} (Notes S1).

### Examples of resilient coding in the {photosynthesis} package for R

We built a package containing analytical tools for plant ecophysiology (Stinziano *et al.*, 2020), embedding our coding principles into the package itself. The R package contains functions for fitting photosynthetic CO_2_ (Farquhar *et al.*, 1980; von Caemmerer, 2000; Gu *et al.*, 2010; Duursma, 2015) and light response curves (Marshall & Biscoe, 1980), temperature responses of biological processes (Arrhenius, 1915; Medlyn *et al.*, 2002; Kruse *et al.*, 2006; Heskel *et al.*, 2016; Liang *et al.*, 2018), light respiration (Kok, 1956; Laisk, 1977; Yin et al., 2009, 2011; Walker & Ort, 2016), mesophyll conductance (Harley *et al.*, 1992), stomatal conductance models (Ball *et al.*, 1987; Leuning, 1995; Medlyn *et al.*, 2011), pressure-volume curves (Tyree & Hammel, 1972; Koide *et al*., 2000; Sack *et al*., 2003), hydraulic vulnerability curves (Pammenter & van der Willigen, 1998; Ogle *et al.*, 2009), and sensitivity analyses (Table 1; Table S2). It also contains functions for modeling C_3_ photosynthesis using the Farquhar-von Caemmerer-Berry biochemical model (Farquhar *et al.*, 1980). The default kinetic parameters for gas exchange fitting procedures are taken from *Nicotiana tabacum* (Bernacchi *et al.*, 2001, 2002). The {photosynthesis} package is currently limited to C_3_ photosynthesis, but future releases should expand its functionality to other photosynthetic pathways. A comprehensive illustration of how to use the package can be found in the vignette of the package (**Notes S2**, “photosynthesis-curve-fitting-sensitivity-analyses.rmd”). There are currently two vignettes available for the package that function as tutorials on CRAN (https://CRAN.R-project.org/package=photosynthesis). The first vignette (titled “photosynthesis-curve-fitting-sensitivity-analyses”) demonstrates how to use curve fitting and sensitivity tools and the second (titled “introduction to the photosynthesis package”) demonstrates how to simulate photosynthetic rate using the Farquhar-von Caemmerer-Berry C_3_ biochemical model, define leaf and environmental parameters, replace default parameters, and solve for chloroplastic CO_2_ concentrations.

**Table 1.** List of {photosynthesis} functions with applications and descriptions. The documentation for each function describes the estimated or simulated parameters, constants, and other calculated values. Documentation is updated to describe new functionalities as they are added.

| Base Functions |  |  |
| --- | --- | --- |
| Applications | Function | Description |
| Gas Exchange | `fit_aci_response` | Fits A-C<sub>i</sub> curves, provides parameters/graphs |
| Gas Exchange | `fit_aq_response` | Fits A-Q curves, provides parameters/graphs |
| Gas Exchange | `fit_g_mc_variableJ` | Fits *g<sub>mc</sub>*, adds *g<sub>mc</sub>* and dCcdA to dataframe for reliability checking |
| Gas Exchange | `fit_gs_model` | Fits the Ball *et al.* 1987, Leuning 1995, and Medlyn *et al.* 2011 models of stomatal conductance, provides parameters/graphs |
| Hydraulics | `fit_hydra_vuln_curve` | Fits the sigmoidal and Weibull models to hydraulic vulnerability data, provides parameters/graphs |
| Hydraulics | `fit_PV_curve` | Fits pressure volume curves, provides parameters/graphs |
| Gas Exchange | `fit_r_light` | Fits *r<sub>light</sub>* according to the Kok (1956) method, Yin method (Yin *et al.* 2009, 2011), or Walker & Ort (2015) method. |
| Gas Exchange, Biochemistry | `fit_t_response` | Fits an Arrhenius (Arrhenius 1915), Heskell (Heskell *et al.* 2016), Kruse (Kruse *et al.* 2006), Medlyn (Medlyn *et al.* 2002), MMRT (Hobbs *et al.* 2013), and quadratic temperature response models, provides parameters/graphs |
| Modeling | `photo` | Simulates C<sub>3</sub> photosynthesis over a parameter set |
| Modeling | `make_parameters` | A set of functions (e.g. `make_enviropar`, `make_leafpar`) that generates the required inputs for `photo` |
| Meta-functions & Utilities |  |  |
| Application | Function | Description |
| Software modification | `check_dependencies` | Generates HTML with package and function dependencies |
| All components | `compile_data` | Compiles the output from the `fit_many` function |
| All components | `fit_many` | Fits a function many times through a grouping variable |
| All components | `print_graphs` | Prints graphs from a list of graphs |
| All components | `sensitivity_analysis` | Allows up to 2-factor sensitivity analysis of any function |

The package is specifically designed to accommodate new analytical tools and discoveries and be easily maintained by new users. Nonlinear curve fitting procedures use the nlsLM function from {minpack.lm} (Elzhov *et al.*, 2016), which provides a more robust fitting procedure for non-linear functions than the base R `nls` function. Graphical outputs are provided using {ggplot2} (Wickham, 2016). Meta-functions were constructed with the tools provided for generalizing functions and arguments in {rlang} (Henry & Wickham, 2019).

The principles of modularity and functional programming have been used to substantially reduce code interdependencies within the software. For example, the `fitaci` function from {plantecophys} has over 30 function dependencies (Fig. 2a). By applying our principles, we were able to reduce this to just 4 function dependencies (Fig. 2b), by reengineering the fitting procedure and eliminating redundant functions and code. Arguably, fewer dependencies could indicate less modularity, even though each of the components is modular, but fewer dependencies may reduce the number of bugs introduced by revisions in other components.

**Figure 2.**
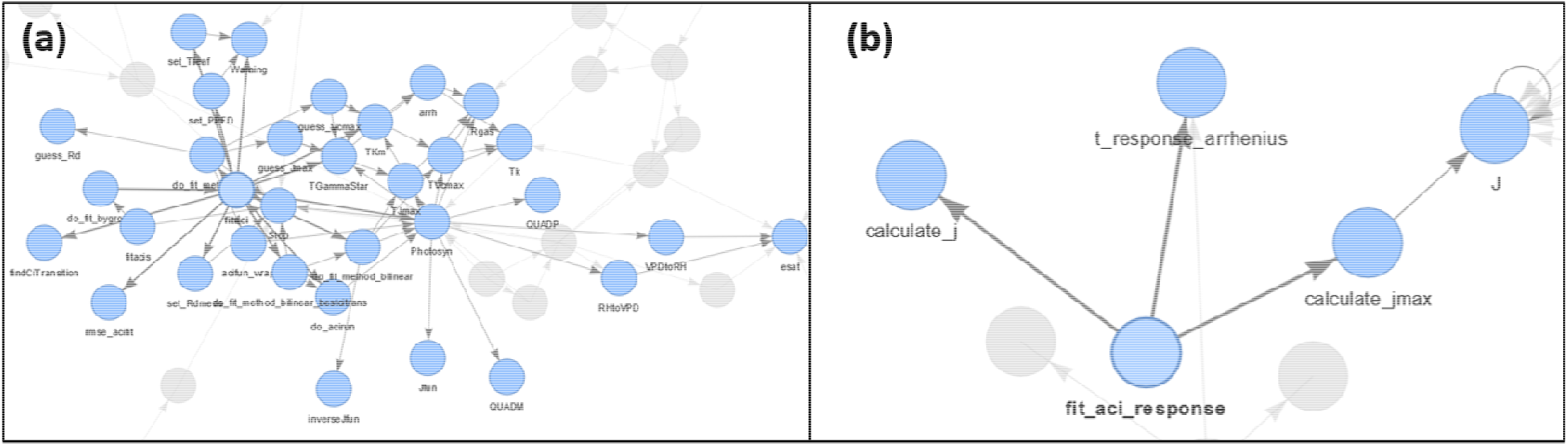
Dependencies of the *A-C_i_* fitting functions in (a) {plantecophys} and (b) {photosynthesis}.

#### Example Dataset

To demonstrate the fitting functions of the package, we use a combination of data collected for the package and previously published data. A CO_2_ by light response curve and CO_2_ by temperature response curve were collected in sunflower (*Helianthus annuum*) grown in a rooftop greenhouse at the University of New Mexico (35.0843°N, 106.6198°W, 1587 m a.s.l., 18.3 to 21.1/15.6 to 21.1 °C day/night temperature with daily irradiances of 600 to 1,200 μmol m^-2^ s^-1^). CO_2_ response curves were measured at irradiances of 1,500, 150, 375, 125, 100, 75, 50, and 25 μmol m^-2^ s^-1^ at a *T_leaf_* of 25 °C. CO_2_ response curves were also measured at *T_leaf_* of 17.5, 20, 22.5, 25, 27.5, 30, 32.5, 35, 37.5, and 40 °C at an irradiance of 1,500 μmol m^-2^ s^-1^. Data to demonstrate hydraulic vulnerability curve fitting methods were drawn from (Hudson *et al.* 2018), while data for leaf pressure/volume analysis come from an unpublished dataset collected at the University of New Mexico. Below we illustrate some of the functionality of the package. This data is freely available in the package so potential users can test out the functions and different analyses in the code. We refer potential users to the package vignette for more worked examples (**Notes S2**, “photosynthesis-curve-fitting-sensitivity-analyses.rmd”).

#### Photosynthetic light response curve fitting

The `fit_aq_response` function returns a list containing the fitted light response model, model parameters, and a graph showing the model fit to the data (Fig. 3a). This function estimates the light-saturated net CO_2_ assimilation rate, quantum yield of CO_2_ assimilation, an empirical an empirical curvature factor, and respiration (Marshall & Biscoe, 1980).

**Figure 3.**
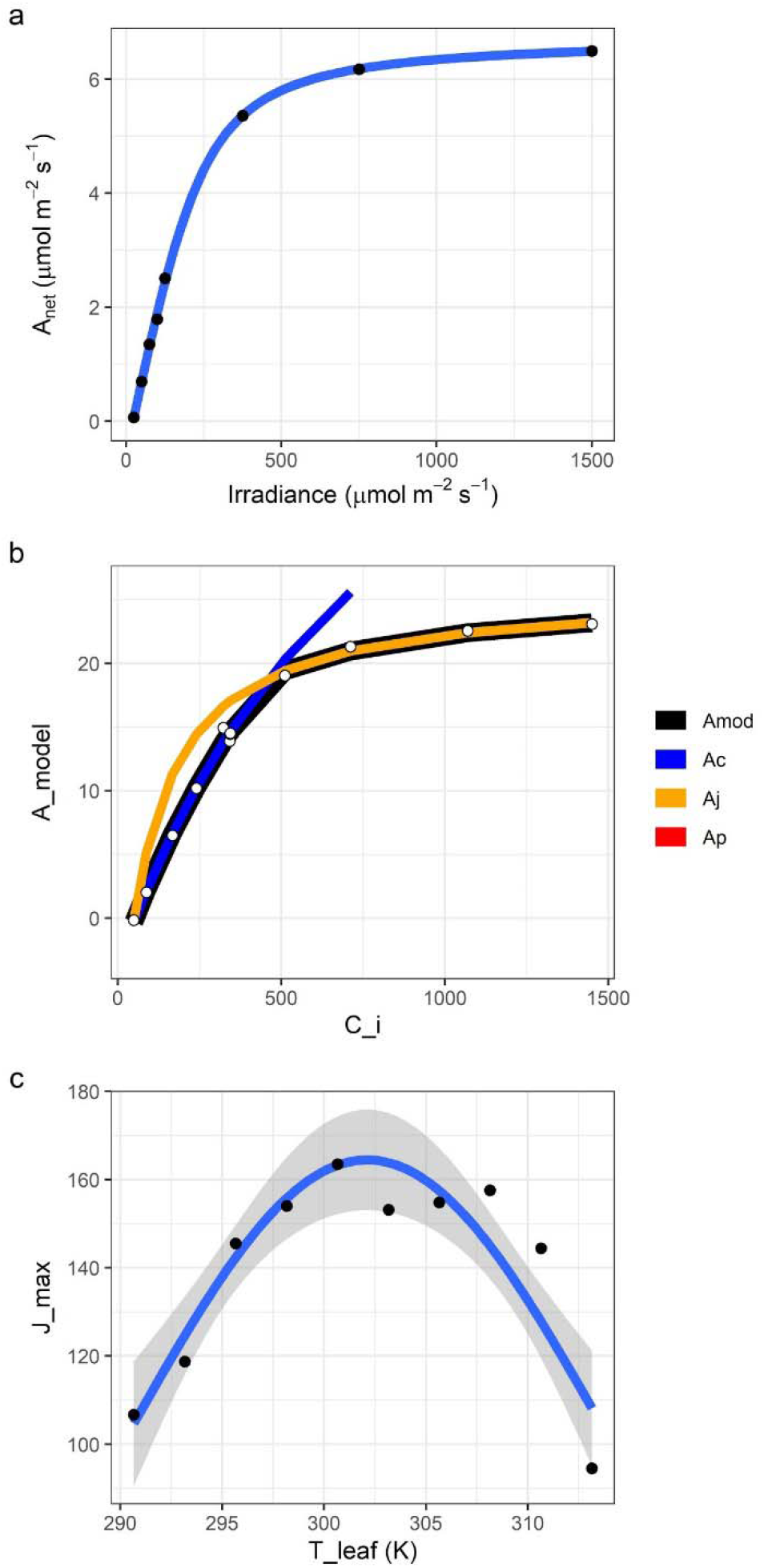
Gas exchange curve fitting outputs. a) Output from {fit_aq_response showing the data (black points), the model fit (blue line), and the standard error on the model fit (grey region). The light response at a [CO_2_] of 100 μmol mol^-1^ is shown. *A_net_:* net CO_2_ assimilation. b) Graph from fit_aci_response showing modelled *A_net_ (A_mod_,* black line), CO_2_-limited *A_net_ (A_c_,* blue), RuBP regeneration-limited *A_net_ (Aj,* orange), triose phosphate utilization-limited *A_net_* (*A_p_*), and the data (white dots). *A_net_:* net CO_2_ assimilation; C_i_: intercellular CO_2_ concentration. c) Output from fit_t_response showing the Heskel temperature response of *J_max_.* Data are black dots, model fit is the blue line, and the grey shaded region is the standard error on the model fit. *J_max_:* maximum rate of electron transport; *T_leaf_:* leaf temperature.

#### Photosynthetic CO_2_ response curve-fitting

The `fit_aci_response` function returns a list containing the fitted parameters, a data frame with the modelled data output, and a graph showing the model fit to the data (Fig. 3b). It estimates the standard parameters of the Farquhar-von Caemmerer-Berry C_3_ biochemical model (Farquhar *et al.* 1980) and parameter standard errors to help evaluate results. As with any nonlinear regression, failure of the solver to converge on a solution or very large standard errors usually indicate problems fitting the model to the data and unreliable parameter estimates.

#### Photosynthetic temperature response curve fitting

A series of temperature response functions can be fit using the package, with the outputs including the fitted model, model parameters, and a graph (Fig. 3c). As with other functions, details about parameters given in the package documentation.

#### Fitting g_m_ using the variable J method

The `fit_g_mc_variableJ` function using the method of Harley *et al.* (1992) using chlorophyll fluorescence and gas exchange data to estimate *g_mc_.* Both *g_mc_* and *δC_c_/δA* are calculated, where *δC_c_/δA* between 10 and 50 are deemed to be “reliable” (Harley *et al.*, 1992), and an average *g_mc_* value is estimated based on the reliable values. This makes it relatively easy to assess the reliability of *g_mc_* estimates (Fig. 4).

**Figure 4.**
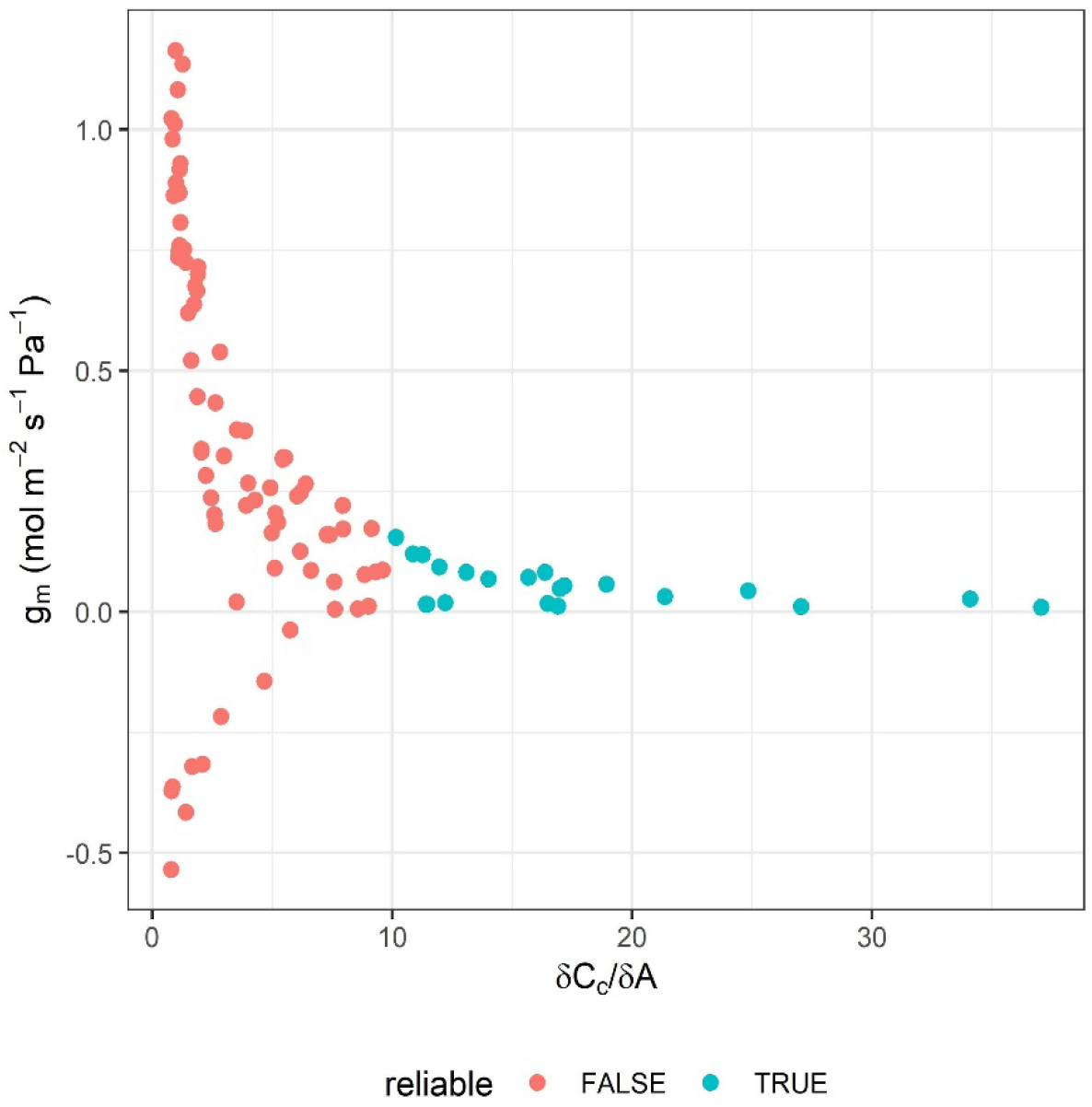
Relationship between *g_mc_* estimated through the variable *J* method and *δC_c_/δA* to test for reliability. The fit_g_mc_variableJ function was used on the CO_2_ by light response data in sunflower. *g_m_*: mesophyll conductance; *Cc:* chloroplastic CO_2_ concentration; *A*: net CO_2_ assimilation.

#### Hydraulic vulnerability curve fitting

The `fit_hydra_vuln_curve` fits hydraulic vulnerability data using both a sigmoidal and Weibull function. Outputs include model fits, parameters and a graph (Fig. 5a).

**Figure 5.**
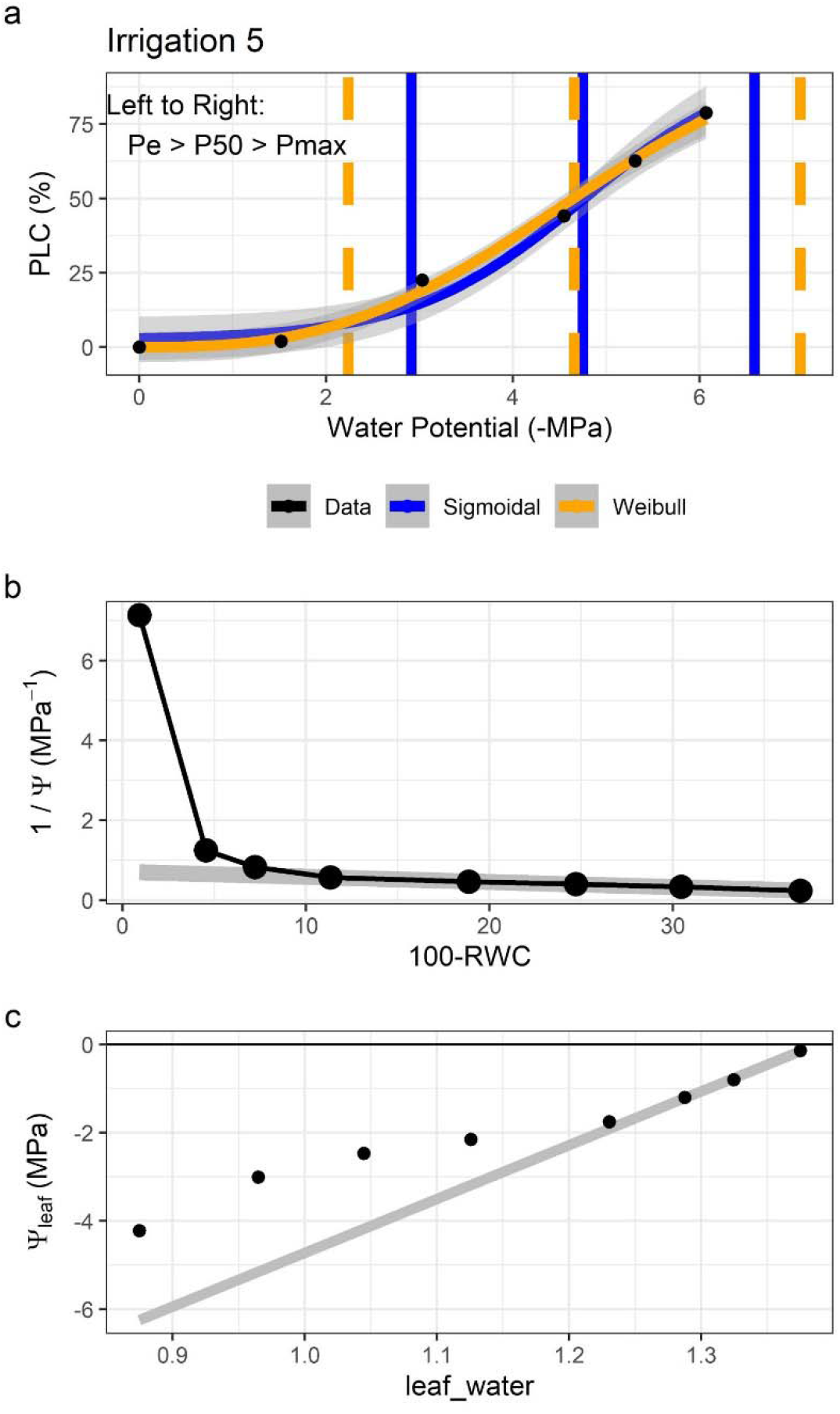
a) Example output from fit_hydra_vuln_curve showing both model fits overlaid on the data (black dots). *PLC:* percent loss of conductivity; *Pe:* air entry point; *P50:* water potential at 50% *PLC; Pmax:* hydraulic failure threshold. b, c) Example output from fit_pv_curve showing the b) water mass graph and c) the pressure-volume curve. Grey lines are fit to the linear regions of the data. Ψ: water potential; RWC: relative water content.

#### Pressure-volume curves

The fit_pv_curve fits pressure-volume curves, returning parameters such as relative water content and water potential at turgor loss points, relative capacitance at full turgor, and others. Outputs include parameters and graphs (Fig. 5bc).

#### Sensitivity analyses

Both `analyze_sensitivity` and `compute_sensitivity` are used in combination for sensitivity analyses. `analyze_sensitivity` allows up to two assumed parameters to be varied in a fitting function, while `compute_sensitivity` runs two types of local sensitivity calculations based on a user-defined reference value: parameter effect (Bauerle et al., 2014) and control coefficient (Capaldo & Pandis, 1997). We can look at the impact of varying *g_m_* and *Γ\** at 25 °C on fitted *V_cmax_* (Fig. 6). We can see that *g_m_* and *Γ\** at 25 °C have an orthogonal impact on *V_cmax_,* with *Γ\** having a stronger control than *g_m_* on *V_cmax_.*

**Figure 6.**
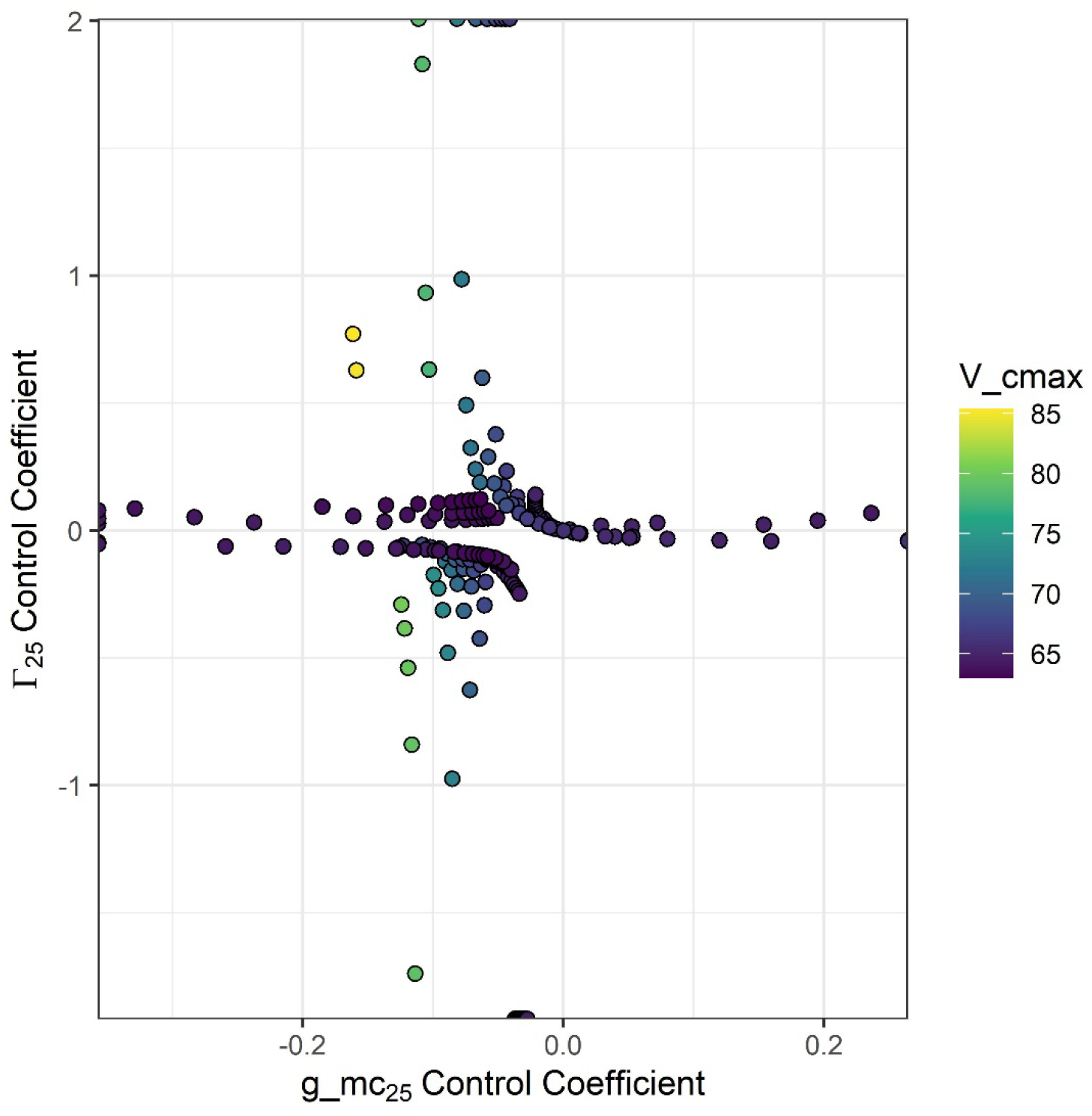
Control coefficients of *g_m_* and *Γ\** at 25 °C calculated from analyze_sensitivity and compute_sensitivity.

### Moving forward - standardized practices and code editors

It is not easy to rewrite software, and we are not arguing as such. Rather, going forward as a community, we argue that we should adopt a set of coding principles and guidelines to create code as flexible as the biology we study. We present the R package, {photosynthesis}, as an example of these principles and guidelines. The consequences of this are not to be understated: it will be easier for new trainees and beginner coders to learn, understand, and write code for the community; and it will be easier to tailor existing code to our projects.

The drawback is that code may run more slowly, which may be a worthwhile tradeoff for some but not others. For example, computational speed may take precedence over flexibility for eddy flux covariance, genomics, and other “big data” applications. In ecophysiology, many datasets are often small enough that even complex analyses may only take 1 hr on one computer core of a multi-core system – as a community we can often afford slower-running code for greater flexibility and ease-of-understanding, especially as this could save days or weeks of coding to write a desired analysis. Our code should be as flexible as, and easier to understand, than the biology it describes.

However, providing code according to these standards is not sufficient - we also need code-competent editorial staff for journals who can properly review and test submitted code to ensure that it runs as intended. In some cases, code for a published dataset does not work even after comprehensive modification (Stinziano, pers. comm.). Standardized coding practices will help to reduce the burden on code editors by making it easier to read and understand code submissions.

## Data Availability Statement

All data and code used in the manuscript are available at https://github.com/cdmuir/photosynthesis.

## Supporting Information

**Methods S1 – Description of variables used in {photosynthesis}**

**Tables S2 – Table of other utility functions in {photosynthesis}.**

**Notes S1 – Benchmark comparison of `plantecophys::fitaci` and `photosynthesis::fit_aci_response`**

**Notes S2 – The {photosynthesis} R package tar.gz file.**

**Notes S3 – Examples tidy data file (hydraulic_vulnerability.csv).**

## Acknowledgements

We thank Tom Buckley, Markus Hauck, and three anonymous reviewers for feedback on earlier versions of the manuscript. This is publication #145 from the School of Life Sciences, University of Hawai’i at Mānoa.

## Supplementary Information

**Table S2.** List of additional {photosynthesis} functions with applications and descriptions.

| Functions |  |  |
| --- | --- | --- |
| Applications | Function | Description |
| Gas Exchange | `aq_response` | Contains the light response model used by `fit_aq_response` |
| Modeling | `bake-par` | Checks that `bake` has worked |
| Modeling | `bake` | Temperature scales input parameters |
| Modeling | `conductance` | Provides calculations for leaf conductances to CO<sub>2</sub> |
| Modeling | `constants` | Checks formatting of physical constant inputs |
| Modeling | `enviro-par` | Checks formatting of environmental inputs |
| Modeling | `FvCB` | Contains the model of Farquhar et al. (1980) for modeling |
| Gas Exchange | `gs_models` | Contains all stomatal conductance models that can be fit with `fit_gs_model` |
| Gas Exchange | `j_calculations` | Contains the electron transport models for fitting $A-C_i$ curves |
| Modeling | `leaf-par` | Ensures proper formatting of leaf parameter inputs |
| Modeling | `parameter_names` | Gets a vector of parameter names |
| Modeling | `photosynthesis` | Contains several functions necessary for `photo` |
| Gas Exchange | `t_functions` | Contains all temperature response functions that can be fit with `fit_t_response` |
| Modeling | `utils` | Contains unit conversion functions for modeling |

